# Strategic exploration in human adaptive control

**DOI:** 10.1101/110486

**Authors:** Eric Schulz, Edgar D. Klenske, Neil R. Bramley, Maarten Speekenbrink

## Abstract

How do people explore in order to gain rewards in uncertain dynamical systems? Within a reinforcement learning paradigm, control normally involves trading off between exploration (i.e. trying out actions in order to gain more knowledge about the system) and exploitation (i.e. using current knowledge of the system to maximize reward). We study a novel control task in which participants must steer a boat on a grid, aiming to follow a path of high reward whilst learning how their actions affect the boat’s position. We find that participants explore strategically yet conservatively, exploring more when mistakes are less costly and practicing actions that will be required later on.

## Introduction

Deciding how to act under uncertainty is a core problem for cognition. Cognitive agents must be able to navigate a world whose dynamics are initially unknown and generally uncertain, learning to generate rewards as they go along. In the context of reinforcement learning, we can think of control as a trade-off between *exploration* (i.e. trying out actions to gain more knowledge about the underlying system) and *exploitation* (i.e. using current knowledge of the system to maximize reward). However, whether and how human explorative control reflects future goals and current uncertainty is still unclear (Wilson, Geana, White, Ludvig, & Cohen, 2014). Are human explorative actions strategic and goal-directed? Or are they rather passive, for instance involving a simple “exploration bonus” that treats uncertainty equally across all actions?

Traditionally, reinforcement learning models have addressed exploration rather implicitly, letting the agent learn about the underlying system *en passant* via outcomes produced while high rewards are chased (Rescorla, Wagner et al., 1972). Exploration, according to this definition, is what happens when an agent optimizes noisily. We will refer to this kind of exploration as *passive exploration.*

More recently, exploration has been incorporated into reinforcement learning models more explicitly via an exploration bonus (Schulz, Konstantinidis, & Speekenbrink, 2016; Wu, Schulz, Speekenbrink, Nelson, & Meder, 2017). An exploration bonus assigns additional utility to less explored actions and thereby assumes that the agent values uncertainty equally across all actions. Exploration, according to this definition, is what happens when expectations are inflated by their attached uncertainties. We will refer to this kind of exploration as *agnostic exploration.*

Another line of research tries to redefine exploration as goal-directed behavior (e.g., Thrun, 1992). The idea behind this approach is that not all uncertainty should be treated equally but rather that exploration should be driven by both the current knowledge of the system and the agent’s overall goal. Exploration, according to this definition, is a strategic action. We will refer to this kind of exploration as *strategic exploration.*

Many real-world control scenarios are non-episodic, such that there are no “second chances” and one may be unable to return to known states. In such situations, one must treat exploration strategically and with great caution to avoid accidents (Klenske & Hennig, 2015). Imagine visiting a country with left-hand traffic from a country with right-hand traffic. Strategically exploring in order to learn how to drive on the left side could allow you to make your mistakes on the quiet roads first before hitting the highway. Moreover, as turning right will be more difficult than you are used to, practicing how to turn right is more important than practicing how to turn left and therefore should be exercised more frequently.

In machine learning, problems of planning under uncertainty have been approached via Bayesian reinforcement learning (BRL; Poupart, 2010), which assigns probabilistic beliefs over the dynamics of a system and the costs of states and actions in order to reason about potential changes to beliefs from future observations, and their influence on future decisions (Duff, 2002). BRL provides a useful framework for assessing strategic exploration behavior as we do here. More specifically, we will make use of the duality between reinforcement learning and control, that is tasks in which an agent has to keep a system at a certain state in order to generate rewards (Feldbaum, 1960; Klenske & Hennig, 2015).

In what follows, we will assess how participants exert control within a novel control paradigm. Therein, strategic exploration allows them to produce greater long-term rewards — formally, within a non-episodic, finite-horizon system with initially unknown dynamics. We will build on recent work by Klenske & Hennig (2015) and assess behavior in two tasks: one in which, due to time-varying state costs, exploration can be delayed until it is more opportune; and one in which the learning agent can distinguish between more and less important exploration of directional actions. We first discuss in more detail the task and the three perspectives on exploration in control theory *passive,agnostic* and *strategic exploration.* We then assess qualitative predictions derived from these in two experiments. We find participants’ behavior to be more in line with predictions derived by *strategic exploration.*

## Control task

In a simple computer game, participants have to navigate a boat as it crosses a sea towards regions in which they can earn a higher bonus. The boat moves incrementally from left to right and by changing its current angle of direction (see Figure 1), participants could attempt to steer the boat up or down, so as to remain in calm waters (blue) and avoid perilous rough seas (red). The overall goal of the game was to minimize the “cost” of the voyage while simultaneously learning both how to control their boat and about an underlying position-dependent “current” that drags the boat off course. In some periods, the area of low cost was very narrow, while in other periods, the area was very wide. Analogous to real sailing, participants had to learn to control the boat through experience, by trying different angles and observing the effect on the boat’s position. This exploration is costly when the low-cost region is narrow, whilst exploration is almost “free” when the low-cost region is very wide.

**Figure 1:**
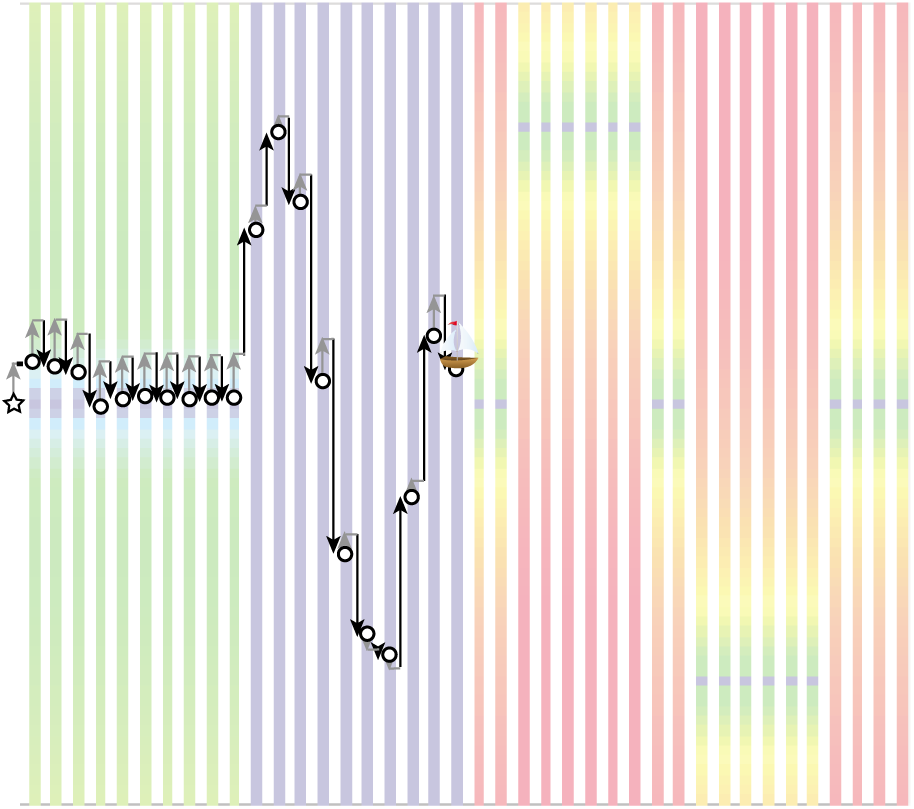
Example path in Experiment 1, Free Late condition. Star = starting position at *t* = 0, circles = subsequent positions. On each trial, the gray arrow shows contribution of underlying current and black arrow contribution of control angle. At *t* = 0 the participant takes a control angle of 0 and drifts upward. On the 10 subsequent trials they attempt to counteract this upward drift by setting a negative angle. During the free exploration stage they use wider angles to explore the variation in the strength of the current at different *y* positions. This allows them to discover that strength of the current is strongest in the center, approaching zero toward the top, constantly pulling the boat upwards.

Our control task is adapted from Klenske & Hennig (2015). Therein, the boat is influenced by two factors, its current position *x*_*1*_ and an underlying current *x*_2_. This means that where the boat will end up on the next trial is influenced by the chosen angle, its current position, and the underlying current which is determined by an unknown nonlinear function. Within our experiments, the underlying current decreased from its full strength in the middle of the sea to zero at the upper and lower edges, and constantly pulled the boat upwards. For example, if participants entered the angle of 0 in the center of the sea, the boat would be pulled upwards more than if they entered the same angle at another position further up. Formally, at each time *t,* the (vertical) position of the boat, *y*_*t*_, depends on a two-dimensional latent state variable *x*_*t*_ and independent random noise *γ*_*t*_ as

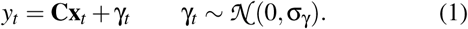

The latent state depends through a nonlinear function on the previous latent state, the controller input (i.e., the chosen angle) *u*_*t*_, and additional *ξ_t_* noise as:

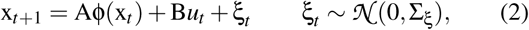

where

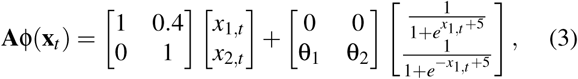

θ = [0.8,0.4]^┬^, and **B** = [0,1]^┬^. The underlying drift is determined by the shifted sigmoid functions on the right-hand side of Equation 3. Given a finite-time horizon with terminal time *T,* the following quadratic cost function was used:

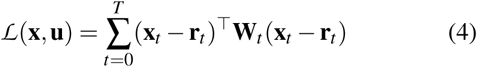

where **r** = [**r**_0_,…, **r**_*T*_] is the target trajectory and **W**_*t*_ the time-varying state cost. The goal of the controller is to find the action sequence **u** = [*u*_*0*_,…,*u*_*T*_] that minimizes the expected cost (and thereby maximizes the expected reward) to the horizon *T.*

## Control strategies

Controlling a system as defined in Eqs. (1) – (3) is difficult, as the state dynamics are nonlinear with an unknown function Φ and parameters **A** and **B**. The controller then not only needs to control the states in accordance to the reference path r, but also learn the parameters (and functions) in order to derive a good control strategy **u**. Thus, the controller not only needs to control the states, but also control her knowledge about the model, hence the term *dual control.*

We will now provide a description of the three different forms of exploration mentioned earlier, and their qualitative predictions in the present control task. The predictions are shown graphically in Figure 2 for the variants of the task used in Experiment 1, which tests whether participants will postpone exploration until it is most opportune, and Experiment 2, which tests whether participants perform strategic (directional) exploration.

**Figure 2:**
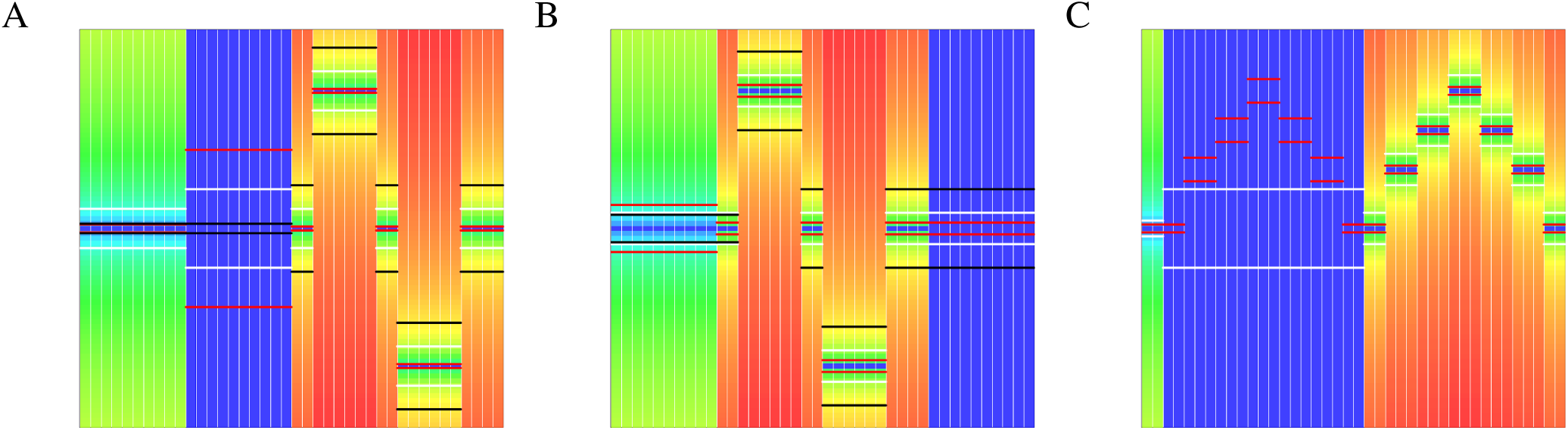
Control environments and qualitative model predictions for Experiment 1 and 2. The agent moves one step right each time point (trials are delimited by white vertical lines) and can control the angle upwards/downwards in which a boat is steering. The background color represent the cost function; the more red, the lower the score; dark blue areas mark free exploration trials. Qualitative model predictions taken from Klenske & Hennig (2015) are represented by horizontal line called ‘predictive regions’. Black lines represent the predictive region for passive exploration, white lines for agnostic exploration, and purple lines for strategic exploration. The more space between the horizontal lines at a trial, the wider the region and the higher the expected variance of the controller’s actions. **A:** Strategic exploration holds off exploration until it comes at lower cost (broader trust region during the dark blue patch) and consequently performs better than passive exploration later on (narrower trust region). **B:** If free exploration phase is moved to the end, strategic exploration explores less overall and expedites exploration to earlier, more costly stages, thereby reducing performance early on in order to achieve the best performance later on. **C:** Instead of agnostically exploring both directions in the same way, strategic exploration uses the free exploration phase to try to move in a trajectory which is rewarding in the future, thereby performing better later on.

### Passive exploration by certainty equivalence

A certainty equivalence controller completely ignores uncertainty about the dynamics and derives a control strategy as if the current (mean) estimates of the system are accurate and knowledge about the system is perfect. Effectively, any learning about the system happens passively, as the control strategy does not focus on minimizing uncertainty. As no active exploration is encoded into this model, it might miss out on important information that could be beneficial to produce better rewards later on. This form of control predicts no exploration, even when exploration is ‘free’ and beneficial to future rewards.

### Agnostic exploration by exploration bonus

To promote exploration, a straightforward adaptation of the certainty equivalent controller is to introduce a Bayesian exploration bonus. Effectively, this means adapting the cost function so that the costs of actions which reduce uncertainty in the model of the control dynamics–as measured by the standard deviation of the posterior distribution over the parameters at each observation point–is temporarily reduced (cf. Srinivas, Krause, Kakade, & Seeger, 2009). This model is still myopic as it only considers uncertainty at the current control step. Moreover, exploration is not strategic, as all uncertainty is treated equally and it does not take into account what knowledge might be most important later on. Under agnostic exploration, the expected behavior would be the attempt to identify all uncertain components, irrespective of their future usefulness.

### Strategic exploration as dual control

BRL involves reasoning about the effect of actions on future rewards and beliefs. Where an exploration bonus renders reducing uncertainty rewarding in itself, in BRL, reducing uncertainty is only attractive insofar as it is expected to result in an increase in future rewards. Optimal BRL requires determining the consequences of strengthening beliefs on future rewards, thereby finding the optimal balance between exploration and exploitation. Unfortunately, the optimal solution to the dual control problem of simultaneously controlling the system as well as possible given current knowledge (exploitation) and learning about the system through experimentation in order to control it better later on (exploration), is known to be intractable.

Approximate dual control involves three conceptual steps which together yield what, from a contemporary perspective, amounts to an approximate solution to Bayesian RL: First, determine the optimal trajectory under the current mean model of the system (as in certainty equivalent control). Second, construct a local quadratic expansion around the nominal trajectory that approximates the effects of future observations. Third, within the current time step *t,* perform the prediction for an arbitrary control input *u*_*t*_ and optimize *u*_*t*_ numerically by repeated computation of steps 1 and 2 at varying *u*_*k*_ to minimize the approximate cost (see Klenske & Hennig, 2015, for implementation). Approximate dual control does not treat all exploration equally but rather explores strategically by, for example, holding off exploration until it is less costly or by exploring actions that will become important later on.

## Experiment 1: Holding off exploration

Our first experiment was designed to test *passive exploration* against both *agnostic* and *strategic exploration* by including a low-cost period which was either introduced relatively early (“Free Early” condition) or at the end of the task (“Free Late” condition). When a low-cost period is introduced early, controllers can make good use of it to explore and better their performance in later periods, while exploring in a low-cost period at the end of the task is not beneficial as there are no later rewards to reap.

Both conditions experienced an initial stage of medium state costs (see Figure 2). However, whereas for the Free Early condition that stage is followed by a stage of free exploration (no costs of errors) which then leads to a stage of very high cost, the Free Late condition experiences the stage with high state costs first before then experiencing the stage with no costs (the two stages are swapped).

We expected participants to behave as strategic controllers and to initially hold off exploration in the Free Early condition until it comes at no cost in the low-cost period, allowing them to be prepared for the most difficult final stages. In contrast, participants in the Free Late condition were expected to explore more in the initial period, in order to be prepared for the second, most difficult stage. In addition, we expected participants in the Free Late condition to explore less in the low-cost period compared to those in the Free Early condition, as late exploration no longer brings benefits if the task is nearly over. Finally, we expected participants in the Free Early condition to generally perform better than participants in the Free Late group, as early exploration would enhance their knowledge of the system for the remainder of the task.

### Design

The manipulation involved changing the order of the reference trajectory (the state values that would produce the highest rewards) and state weightings.

In the Free Early-condition the reference trajectory and state weightings were:

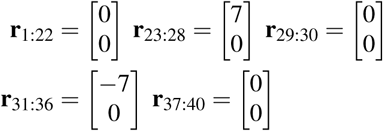

and

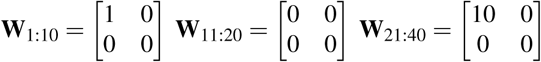

In the Free Late condition, these were:

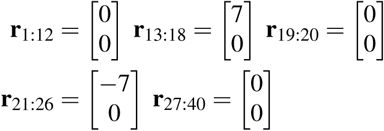

and

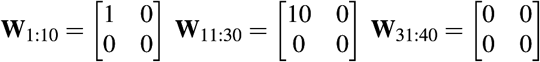

### Materials

Participants were told that they had to navigate a boat through the sea in a sailing competition. On every trial, their boat was at a current position (*y*_*t*_) and they had to determine an angle (*u*_*t*_, between -180° and 180°) in which they wanted to sail. Additionally, they had different target areas (*r*_*t*_) on each trial marked by dark blue colors and how far they were off from the target area was penalized differently (based on **W**_*t*_). An example trial from the task (for the Free Early condition) is depicted in Figure 1.

The cost function was shown to participants through the color of each position in the sea. Participants could earn between 0 (positions with a red background) and 100 points (positions with a blue background) per trial.

### Participants

Sixty-one participants were recruited via Amazon Mechanical Turk and received $1 and a bonus of up to $1. Thirty-nine participants were male and the mean age was 31.3±8.4.

### Results

The distribution of boat position, as well as average chosen angles, are depicted in Figure 3. We can see that, overall, participants managed to steer the boat reasonably well. A linear regression of condition, cost function weights (coded as dummy variables), and trial number onto participants’ scores (see Table 1) showed that, unsurprisingly, cost function weights had the largest effect on participants’ scores. Moreover, performance increased significantly over trials. Importantly, condition affected overall performance, such that participants in the Free Early condition performed better than participants in the Free Late condition. This confirms the hypothesis that participants would benefit from early free exploration.

**Figure 3:**
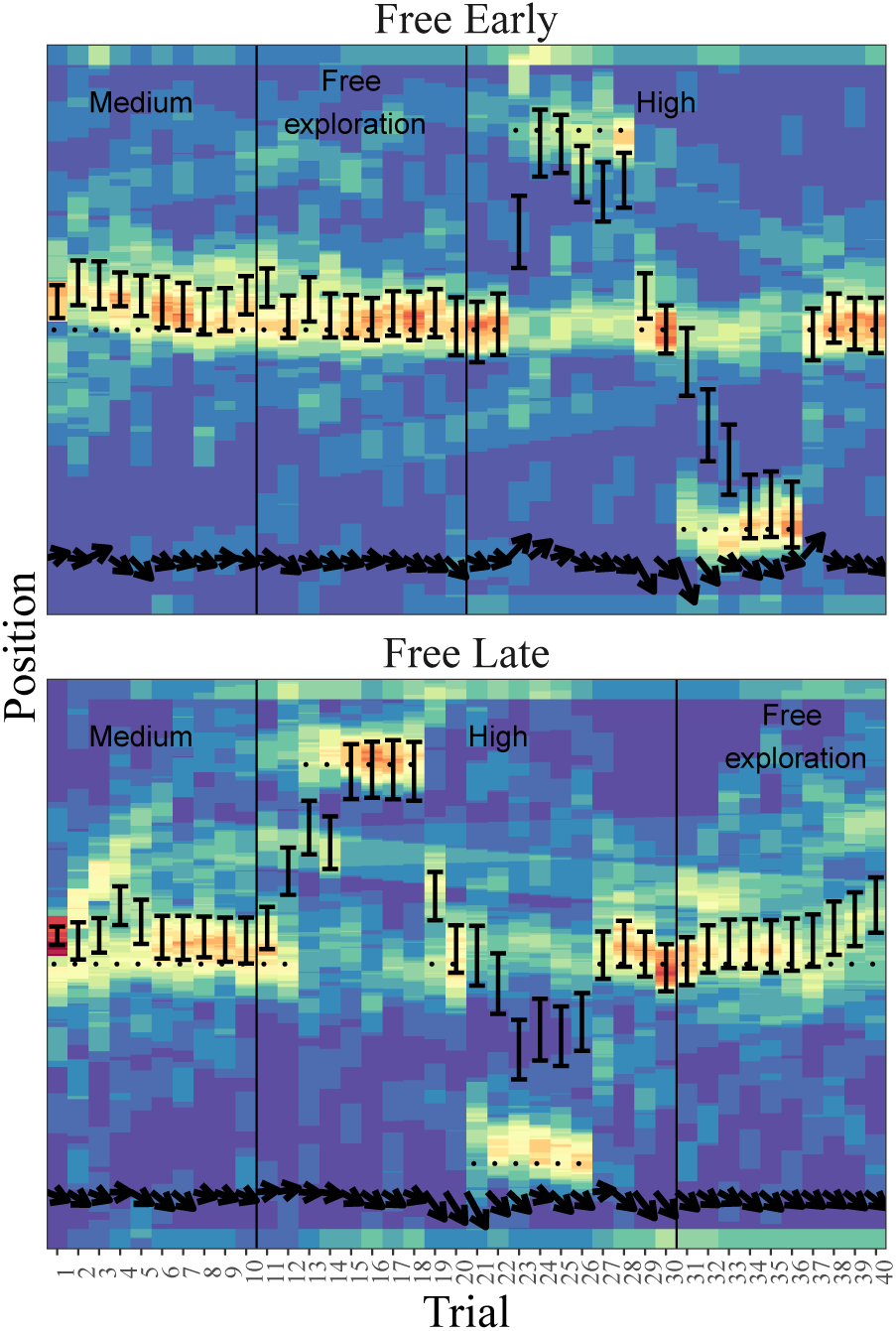
Boat positions by condition in Experiment 1. The heat map reflects number of participants who were at that position on a given trial. Error bars represent the standard error of the average position per trial. Arrows indicate the average chosen angle. Black dots mark the target trajectory and periods with different state weights are delimited by vertical lines.

**Table 1:**
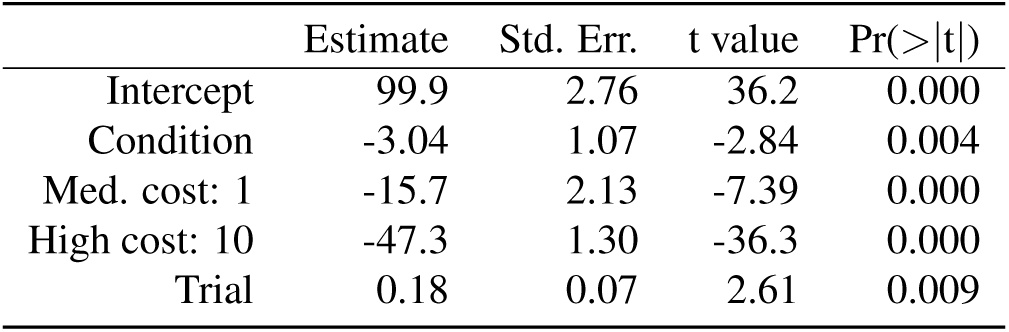
Regression estimates for Experiment 1. *r*^2^ = 0.38.

Another hypothesis was that participants in the Free Early condition would explore more during the free exploration stage than participants in the Free Late condition. Confirming this hypothesis, the participant-wise variance of chosen angles during the free exploration stage was significantly larger for the Free Early condition than for the Free Late condition (*t* (59) = 2.62, *p* < 0.01). As such, participants indeed seemed to strategically adapt their exploration behavior to the underlying cost function.

While we expected participants in the Free Late condition to explore more in the initial stage of medium difficulty than those in the Free Early condition, a similar test to the one above did not confirm this (*t* (59) = 0.63, *p >* 0.5). As such, there is no clear evidence that participants in the Free Late condition used the medium difficulty period to explore in order to perform better in the high-difficulty period.

Overall, participants in Experiment 1 showed hallmarks of strategic exploration. However, they did not explore as vigorously as approximate dual control predicted, often only doing so during completely free exploration periods. As soon as exploration is somewhat costly, participants seem to shift focus back to normal (perhaps certainty-equivalence based) control, thereby more conservatively trading off between exploration and exploitation.

## Experiment 2: Directional exploration

The second experiment was designed to distinguish between *agnostic* and *strategic exploration*, involving the explicit exploration of directional actions. The design was again based on ideas put forward by Klenske & Hennig (2015). In both conditions, a free exploration phase was followed by a high difficulty period, in which controllers either had to move the boat first up then down again (Up-Down condition) or first down and then up again (Down-Up condition).

If exploration is indeed strategic rather than agnostic and simply based on an exploration bonus, then participants in the Up-Down condition should focus exploration in the free exploration phase on first learning to travel precise increments upwards and then precise increments downwards, whereas participants in the Down-Up condition should explore to do the opposite, as knowledge about these actions will be useful later on. Note that mimicking the later target trajectory during the free exploration phase is better then trying upwards and downwards movements at one position as the current, and with that the effect of a chosen angle on the position, varies nonlinearly depending on the boat’s position.

### Design

The underlying dynamics were exactly the same as in Experiment 1. The manipulation solely concerned the reference trajectory, which for the Up-Down condition was:

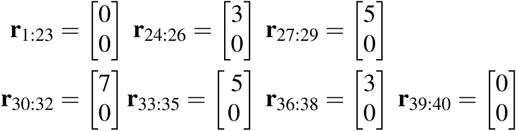

And for the Down-Up condition, the reference trajectory was:

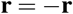

The state weighting was the same for both groups:

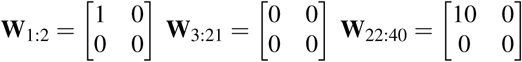

### Materials

Participants were again told that they were taking part in a sailing contest. Participants in the Up-Down condition were then shown the control environment sketched out in Figure 2 (right panel), whereas participants in the Down-Up condition experienced the same control environment but flipped around the center horizontal axis.

### Participants

Forty-six participants were recruited via Amazon Mechanical Turk and received $1 and a bonus of up to $1. 16 participants were female and the mean age was 34.32±11.17.

### Results

Figure 4 shows participants’ boat position by group. Again, participants seemed to be able to learn how to steer the boat towards its targets in both groups. As before, we performed a linear regression of the weights, trials and condition onto participants’ score (see Table 2).

**Figure 4:**
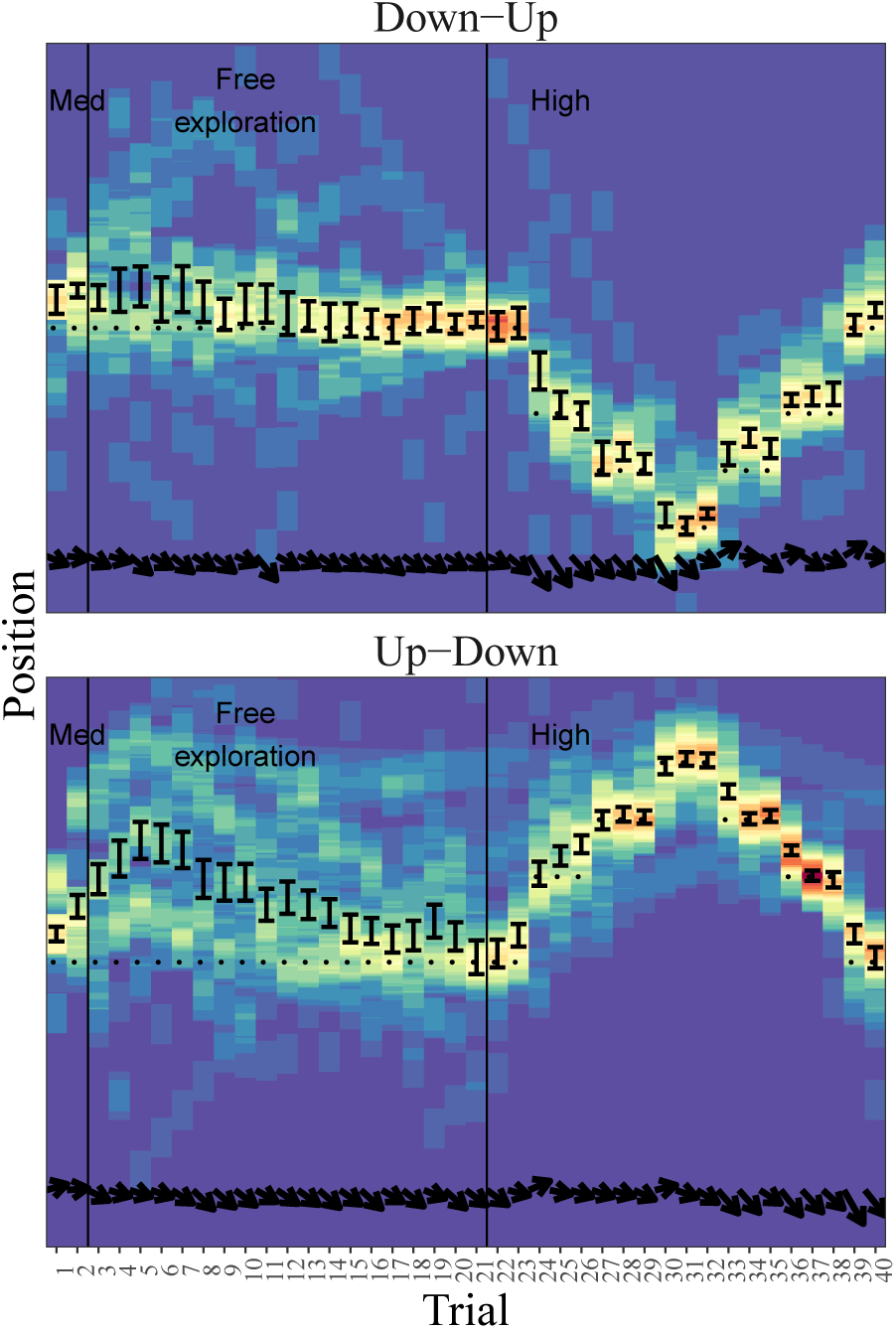
Boat positions by condition for Experiment 2. See legend of Figure 3 for further details.

**Table 2:** Regression estimates for Experiment 2. *r*^2^ = 0.45.

|  | Estimate | Std. Err. | t-value | Pr(>\|t\|) |
| --- | --- | --- | --- | --- |
| Intercept | 97.8 | 2.72 | 35.9 | 0.000 |
| Condition | 0.23 | 1.75 | 0.13 | 0.89 |
| Med. cost: 1 | -11.3 | 1.69 | -6.56 | 0.000 |
| High cost: 10 | -26.6 | 1.37 | -19.4 | 0.000 |
| Trial | 0.16 | 0.06 | 2.41 | 0.01 |

The weights had again the largest effect on participants’ scores and participants’ scores improved over time. There was no significant difference between the scores of the two conditions.

Strategic exploration is visible in the Up-Down condition as participants’ mean position goes up and then down again, thereby showing clear signs of practicing the route to come. This can also be found by testing the difference between condition’s average position during times of free exploration, which was significantly higher for the Up-Down condition (*t*(44) = 3.21, *p* < 0.01). Strategic exploration was not as pronounced in the Down-Up condition, as the mean position seems closer to a straight line than the later target trajectory. Since the prevailing current would nudge any passive participants who aimed straight ahead upward, a bias toward the upper half is to be expected in both conditions. There is no evidence that participants in the Down-Up condition explored less, as there was no difference in the variance of chosen inputs during the free exploration phase between the conditions (*t* (44) = −0.32, *p* > 0.75). Participants in the Down-Up condition chose angles which were on average more downwards during the first 10 trials (*t* (44) = −3.17, *p* < 0.01). Thus, perhaps participants in the Down-Up condition also explored strategically, but were less able to steer the boat in a clear and consistent “practice run” of the desired future route.

## Discussion and Conclusion

Scenarios in which we have to explore to effectively exploit dynamical systems are ubiquitous in daily life. We introduced a novel control task and assessed to what extent people’s exploration can be seen as a strategic, opportunistic, and goal-directed behavior.

We found that participants displayed hallmarks of strategic exploration, exploring differently depending on the cost function and, in some cases, practicing part trajectories which would become important later on. However, strategic exploration seemed more conservative than that of an idealized approximate dual control strategy. During periods of medium cost, participants seemed reluctant to explore in order to benefit their performance during a following high-cost period in Experiment 1. For controllers who learn and choose actions more noisily than statistical algorithms, perhaps the future benefits of this costly exploration did not outweigh the immediate costs. Participants also did not always play out strategies of future importance during free exploration trials as indicated by Experiment 2. As participants in the Up-Down condition could easily follow the underlying upward-current, participants in the Down-Up condition had to go against the current in order to explore strategically. Therefore, the difference in exploration behavior could imply that, for humans, serendipity still plays a part in discovery of effective exploration strategies.

As strategic exploration requires considerable planning, even when dual control is approximate, it is likely to require considerable mental effort. Future research could look into possible heuristics which approximate strategic exploration whilst further reducing computational costs.

